# Thermodynamic Characterization of LC3/GABARAP Ligands by Isothermal Titration Calorimetry

**DOI:** 10.1101/2024.01.08.574610

**Authors:** Johannes Dopfer, Martin P. Schwalm, Stefan Knapp, Vladimir V. Rogov

## Abstract

Isothermal titration calorimetry (ITC) is a widely used technique for the characterization of protein-protein and protein-ligand interactions. It provides information on the stoichiometry, affinity, and the thermodynamic driving forces of interactions. This chapter exemplifies the use of ITC to investigate interactions between human autophagy modifiers (LC3/GABARAP proteins) and their interaction partners, the LIR motif containing sequences. The purpose of this report is to present a detailed protocol for the production of LC3/GABARAP-interacting LIR peptides using *E. coli* expression systems. In addition, we outline the design of ITC experiments using the LC3/GABARAP:peptide interactions as an example. Comprehensive troubleshooting notes are provided to facilitate the adaptation of these protocols to different ligand-receptor systems. The methodology outlined for studying protein-ligand interactions will help to avoid common errors and misinterpretations of experimental results.

## 1 Introduction

Autophagy is an essential cellular process for maintaining homeostasis by removing various unwanted cellular components (autophagy cargo) [1]. Different types of cargo, such as proteins and their aggregates, organelles, pathogens, and lipid droplets, are targeted for lysosomal degradation by autophagic pathways [2]. Macroautophagy, hereafter simply referred to as autophagy, employs double-membrane vesicles called autophagosomes, as carriers to transport autophagosome-encapsulated cargo to lysosomes for degradation. An large number of autophagy-related (Atg) proteins coordinate this process [3]. The human homologs of the yeast Atg8 protein (termed LC3/GABARAP proteins) play a critical role within the selective autophagy machinery by coordinating the autophagosome biogenesis and tethering the autophagy receptors to the forming phagophore through multiple protein-protein interactions [4].

Recent studies have shown that the autophagy receptors (and other LC3/GABARAP-interacting proteins) contain a short sequence motif called the LC3 interacting region (LIR). This motif is the recognition sequence that mediates a variety of Atg8-related protein-protein interactions [5]. The LC3/GABARAP proteins have an LIR-docking site (LDS) that is recognized by canonical LIR motifs with moderate affinities. Variations and extensions of the canonical LIR modulate its affinity for LC3/GABARAP, resulting in binding affinities ranging from subnanomolar to millimolar [6]. An additional binding site on the surface of LC3/GABARAP proteins has been reported called the UIM-docking site [7]. This site has not been extensively characterized biochemically or in terms of sequence requirements present in interaction partners. Investigation of interactions between LC3/GABARAP proteins and LIR motifs on affinity and selectivity remains in the focus of the contemporary field of autophagy. Interactions mediated by LIR binding site provides the basis for the design of autophagosome-tethering compounds (ATTECs [8, 9]) which may recruit protein of therapeutic interest to the autophagy degradation machinery [10]. To elucidate the biophysical nature of the key LC3/GABARAP:ligand interactions a comprehensive analysis of structural and thermodynamic factors driving these interactions is required.

Calorimetric measurements determine the amount of heat released in an exothermic reaction or consumed in an endothermic reaction. The measurement method used in isothermal titration calorimetry (ITC) is based on the power compensation required to maintain a constant temperature difference between two measurement cells. ITC is a general method for studying chemical reactions and molecular interactions based on a characteristic change of enthalpy. To study LC3/GABARAP:peptide or LC3/GABARAP:small molecule interactions, ITC marks a valuable, label-free method. A titration calorimeter consists of a sample cell and a reference cell enclosed in an adiabatic shield. To ensure isothermal conditions, the temperature difference between the reference cell is continuously monitored and adjusted by a heating module. When a ligand is titrated into a protein solution in the sample cell, either an endothermic or exothermic reaction will be initiated by the protein-ligand interaction. The resulting temperature difference between the sample and the reference cell is recorded using a Peltier device located between the two cells (Fig. 1). A feedback system regulates the power compensation required to maintain both cells at the defined temperature, which is recorded over time, and converted into a readout of µcal/s or µJ/s, often referred to as a thermogram [11, 12]. Integrating the injection peaks over time yields the energy required to return back to the baseline value. A binding isotherm is calculated by plotting the heat of each injection normalised to the amount of injected ligand against the molar ratio. By using a nonlinear fitting model, the association constant *K*_*a*_, the total change in binding enthalpy Δ*H* and the stoichiometry (*n*) of the monitored interaction can be calculated. The simplest binding model assumes a single binding site describing the binding of a ligand (L) to a macromolecule (M). As described by the law of mass action, *K*_*a*_ is related to the activity of the macromolecule-ligand complex and its constituents by:

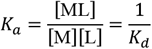

The dissociation constant *K*_*d*_ is inversely related to *K*_*a*_. The Gibbs free energy of binding ΔG is linked to *K*_*a*_ by the following equation:

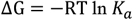

with the universal gas constant R and the absolute temperature T. ΔG may also be expressed in terms of Δ*H* and Δ*S* as:

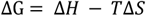

Thus, a binding event consists of two distinct energy-related terms, both of which must be considered in order to understand its underlying intricacies [13]. A properly designed ITC experiment is therefore capable of determining the total change of enthalpy Δ*H*, the association constant *K*_*a*_, the stoichiometry coefficient *n* and the Gibbs free energy ΔG of the process, allowing the calculation of the change in entropy Δ*S* at a certain temperature and buffer condition.

**Figure 1.**
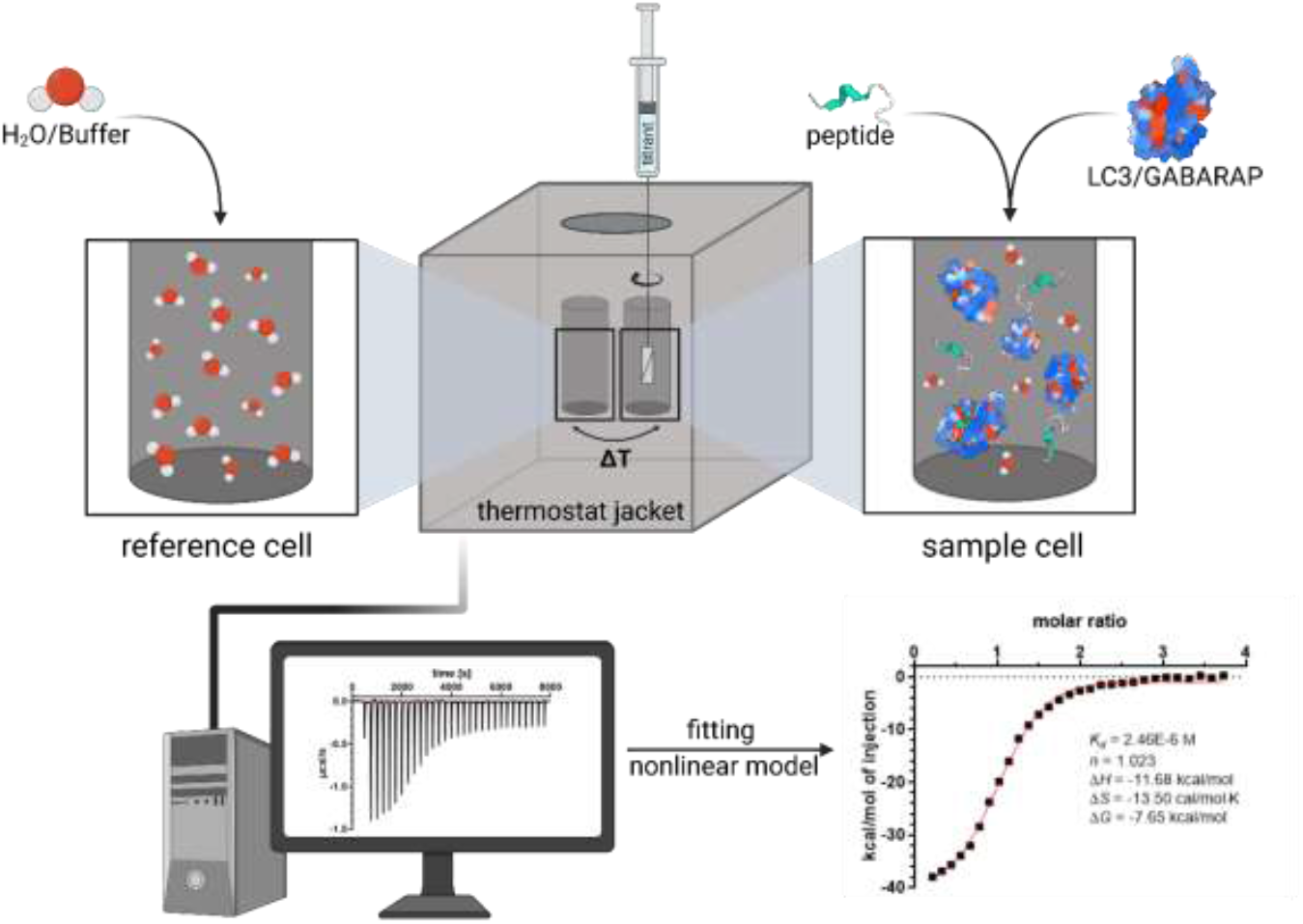
Schematic overview of a titration calorimeter consisting of a sample cell and a reference cell. The titrant is injected into the sample cell and mixed to ensure rapid diffusion. As a result of binding events, there is a difference in the temperature which is detected and transferred into a feedback system to maintain isothermal conditions. The recorded differential power signal is integrated for every injection peak and analyzed by fitting a nonlinear binding model.

The shape of the binding isotherm depends on *K*_*a*_, the total concentration of the macromolecule [M]_*t*_ (if placed as receptor in the sample cell) and the stoichiometry *n* of the binding interaction [14]. The parameter describing this relationship is termed *c* value and it can be expressed in terms of the *K*_*a*_ or *K*_*d*_:

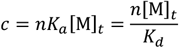

Ideally, for accurate calculation of the thermodynamic parameters mentioned above, the value of *c* should between 10 and 500. A low *c* value titration (*c* < 5) has to be performed if the solubility of M acts as limiting factor for the adjustment of [M]_*t*_ or low affinity ligands (in the micromolar range) are expected. Experiments conducted under these conditions can still lead to accurate determination of *K*_*a*_ and consequently Δ*G*, if the protein saturation is sufficient, the concentrations of both ligand and protein are well known, and the stoichiometry is fixed to the actual value. Caution should be taken by interpreting the value of Δ*H* in case of low *c* value titrations [15]. For *c* values larger than 1000, the slope of the sigmoidal binding isotherm becomes exceedingly steep, closely resembling a step function. Under high *c* value conditions, the accurate determination of *K*_*a*_ is not possible, while the measured enthalpy change of the binding event is still reliable. Lowering [M]_*t*_ to reduce *c* is limited by the sensitivity of the calorimeter. Therefore, one method for determining high affinity binding equilibria relies on artificially reducing the binding affinity by altering the experimental conditions. For instance, temperature modification can be implemented in combination with the determination of thermodynamic linkage parameters. This approach permits the estimation of the Gibbs free energy change of the altered condition and its extrapolation back to the condition of interest. In case of the change in temperature this can be achieved by utilizing a modified van’t Hoff equation and recorded values of the heat capacity changes for different temperature settings [16]. It is important to exercise caution, as extrapolation can introduce errors and changes in experimental conditions may alter the stability of the system being studied, which could ultimately result in protein aggregation. Another approach for studying high affinity ligands (*K*_*d*_ < 1 nM) is a displacement titration. The method involves saturating the receptor with a weaker competitive ligand before titrating the high affinity ligand, resulting in a reduced apparent affinity [17].This is due to an energy penalty which is paid for displacing the weaker ligand from the binding site, increasing the Gibbs free energy change of the high affinity binding event. For this experimental mode at least two titrations are necessary: I) the association constant of the lower affinity ligand *K*_*low*_ is to be determined II) displacement of the weaker ligand by titration of the high affinity ligand to yield its apparent association constant 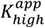. Assuming *n* identical and independent binding sites, fitting a binding isotherm to the data given the additional information of *K*_*low*_ and the concentration of the weaker ligand [X] is possible [18]. The following equations describe the relation between the association constants and the change in enthalpy:

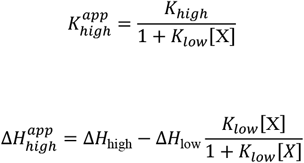

This technique has been utilized by Li et al. to study the low nanomolar interactions between the LIR of ankyrins and different GABARAP isoforms [19], and this method has been described by another protocol [17]. The thermodynamic signatures of the binding of extremely weak ligands, such as fragments that may not have high enough affinity to reach saturation without encountering a solubility problem, can also be analyzed using the principle of displacement titrations. In this case the macromolecule is incubated with the fragment of unknown affinity, which is then displaced by a molecule with known affinity that may contain the fragment as a substructure [20].

## 2 Materials

Peptides were either generated as described in this protocol or obtained from GeneCust. Initial references to chemicals and supplies include supplier and catalogue information. The expression plasmids for LC3A, LC3B, LC3C, GABARAP, GABARAPL1, GABARAPL2 and human ANK3 LIR motif can be obtained upon request.

## 3 Methods

### 3.1 Peptide/Protein Expression and Purification

ITC measurements rely on highly purified protein in a well-controlled buffer system. The plasmid designed for ANK3 peptide expression features a T7 promoter and it is suitable for expression in *E. coli* Rosetta (DE3) cells, along with a ubiquitin-based-tag that is cleavable by TEV protease. The solubility-enhancing Ub-leader consists of a modified ubiquitin and a His_10_ sequence designed to increase the yield and allow for purification via His-tag mediated affinity chromatography [21]. The subsequent steps for protein expression and purification were carried out as described:

#### Bacterial Transformation

1. Thaw 50 µl of *E. coli* Rosetta (DE3) cells (Sigma-Aldrich, Cat# 71400-3) on ice and add approximately 1–10 ng of plasmid DNA. Incubate the cells on ice for 20 minutes, followed by a brief heat shock for 30 seconds at 42 °C. After heat shock, incubate the cells for 2–3 minutes on ice before adding SOC medium (provided in the Rosetta kit). Alternatively, LB (Carl Roth, Cat# X969.1) can be used. Incubate the cells for one hour at 37 °C while shaking at 300 rpm. To concentrate the transformed cells, harvest the cells at 5000 g for five minutes and remove the supernatant. Subsequently, 50 µl of fresh medium is added and the cells are plated on LB-agar plates (Carl Roth, Cat# X969.1) containing 50 µg/ml Kanamycin (Carl Roth, Cat# T832.4) and 34 µg/ml Chloramphenicol (Carl Roth, Cat#3886.2). Incubate the plates at 37 °C overnight.
2. The following day, use a sterile pipette tip to inoculate 100 ml TB culture medium (Carl Roth, Cat# 3556.2) containing 50 µg/ml Kanamycin and 34 µg/ml Chloramphenicol, followed by incubation overnight at 37 °C while shaking at 180 rpm.

#### Protein Expression

3. For each construct isolation, prepare a total of 4 L of sterile TB medium supplemented with 50 µg/ml Kanamycin distributed across four 2 L shaking flasks (1 L TB medium per flask).
4. Inoculate each of the expression cultures with 25 ml of the pre-culture, followed by incubation at 37 °C while shaking at 120 rpm to an optical density at 600 nm (OD_600_) of 0.8.
5. Before inducing protein expression, cool the cultures to 18 °C and shake them for an additional 30 minutes at 120 rpm. Use 1 mL of a 0.5 M a stock solution of Isopropyl-ß-D-thiogalactopyranoside/IPTG (Carl Roth, Cat# CN08.1) per flask and continue shaking overnight at 18 °C and 120 rpm.
6. Harvest the cells by centrifugation of the cultures for 10 minutes at 4000 g and discard the supernatant. Transfer the cell pellet in a 50 ml tube (Greiner Bio, Cat# 227270), which can be stored at -20 °C for future purification. Keep the cells on ice, if purification is continued.
7. For lysis, resuspend the cells in a lysis buffer containing 500 mM NaCl (Carl Roth, Cat# 3957.1), 0.5 mM Tris-(2-carboxyethyl)-phosphine Hydrochloride/TCEP (Carl Roth, Cat# HN95.2), 30 mM 4-(2-Hydroxyethyl)piperazine-1-ethanesulfonic acid/HEPES at pH 7.5 and 5% glycerol (Carl Roth, Cat# 3783.1), before the addition of 1 tablet of Complete protease inhibitor (Sigma Aldrich, Cat# 11836145001) and a few flakes of lyophylized DNase I (Sigma Aldrich, Cat# 4716728001) (alternatively polyethylenimine can be used for DNA precipitation).
8. Sonicate the cell suspension on ice for 15 minutes, using an alternating 5-seconds on and 10-seconds off cycle. After mixing the lysate repeat sonication for another 15 minutes.
9. Separate the lysate from cellular debris by centrifugation at 4 °C and 50000 *g* for 1 h followed by filtration, using a 0.45 µm sterile filter (Carl Roth, Cat# XA50.1).

#### Protein Purification

10. The protein within the filtered lysate is subsequently isolated, utilizing a chromatography system (e.g. Äkta Start, Cytiva; alternatively, gravity flow columns can be used) to load the lysate onto a Ni-NTA affinity chromatography column (Cytiva, Cat# 17524801). After loading, wash the column with 3 column volumes (CV) of lysis buffer containing 20 mM Imidazole (Carl Roth Cat# X998.1). For elution, change the buffer to lysis buffer containing 300 mM Imidazole in a stepwise manner and monitor the A280 nm chromatogram while fractionating.
11. Perform a SDS PAGE gel electrophoretic analysis of the cleared lysate (collected before affinity chromatography), the 20 mM imidazole wash fraction, and all elution fractions corresponding to a signal at A280 nm (include a Protein marker for protein size estimation e.g., CozyHi™ Prestained Protein Ladder, HighQu Cat# PRL0202). After running the SDS PAGE at 200 V for 1 h, stain the gel using stains such as e.g. ROTI®Blue quick (Carl Roth, Cat# 4829.1) to visualize the protein bands corresponding to the purified Ub-fused protein and assess the preliminary purity.
12. To cleave the His_10_-tag from the purified protein, pool the fractions with corresponding protein size (while avoiding impure fractions) according to the SDS PAGE gel and prepare a sample for the subsequent SDS-PAGE. Add approximately 500 µg purified His-tagged TEV protease (Sigma Aldrich, Cat# T4455) to the pooled fractions and transfer the mixture into a 50 ml reaction tube. Incubate the tube on a tube roller mixer (Carl Roth, Cat# XK30.1) at 4 °C overnight.
13. Prepare another SDS-PAGE gel by loading the sample collected at step 12 before cleavage and a sample collected from the reaction tube after incubation with TEV protease to evaluate the cleavage efficiency. Note, that the cleaved peptide is of ~3 kDa, hence use a SDS PAGE gel of sufficient acrylamide concentration (~20 %) to achieve appropriate resolution. Stain the gel to visualize the bands corresponding to the purified peptide without tag by comparing the size difference between the uncleaved and cleaved fractions.
14. Assuming that the molecular weight of TEV protease is 27 kDa, the uncleaved fusion protein is 15 kDa and Ub-tag is 12 kDa, the cleaved peptide can easily be separated by using a spin concentrator with 10 kDa cut-off (Sigma Aldrich, Cat# UFC901008) and collection of the *flow-through*. Alternatively, size exclusion chromatography (SEC) on the prepacked Superdex 75 PG HighLoad 16/600 column (Cytiva, Cat#28-9893-33) can be used to purify the target peptide as well.
15. To yield adequately concentrated peptide for ITC experiments, use a spin concentrator with 3 kDa pore size, which is sufficient given the larger hydrodynamic radius of the unstructured peptide. This also provides a convenient opportunity to exchange the buffer for the one used in the subsequent ITC experiments.
16. Ultimately, the identity and purity of the peptide can be confirmed through mass spectrometry.

### 3.2 Peptide Measurement

#### 3.2.1 Experimental Design

For the measurements conducted in this work, we employed the standard volume Nano ITC calorimeter from TA Instruments, which is equipped with an active cell volume of 1.0 mL. The volume per injection and stir rate might vary depending on the system (refer to manufactures manual, if a comparable device is used). Throughout the experiment, microaliquots are injected into the sample cell, while the contents of the sample cell are kept in continuous motion by the rotation of the syringe and the paddle at its tip, as depicted in figure 1. The stirring rate should be sufficiently high to allow rapid diffusion of the macromolecule/ligand introduced during the injections. Excessive stirring however, decreases the signal-to-noise ratio due to the formation of artificial heat signals caused by friction. The recommended stirring rate differs between instruments. Here, all experiments were conducted at 350 rpm (recommended 300–400 rpm) and yielded a stable baseline. The injection volume is dependent on the volume of the sample cell and the sensitivity of the calorimeter (5–10 µl for the Nano ITC). If the binding event causes large differences in heat, the volume of the aliquots may be reduced. It is common practice to use a smaller initial injection volume, that is not taken into account when analyzing the data, to exclude the diffusion of the cell contents into the syringe tip during equilibration from the analysis. The spacing of the injections has to be large enough to avoid overlapping of the injection peaks. During our method optimization, the standard 300 s injection spacing was reduced to 250 s to shorten the total duration of the experiment. The number of injections is typically chosen to allow for the collection of enough data points to accurately fit the binding isotherm, while also generating an acceptable heat signal per injection (usually up to 30 injections).

The concentrations of macromolecule and ligand in each experiment depend on the extent of heat change upon binding and the anticipated binding constant. For a 1:1 binding reaction, the molar ratio (ligand to protein) at the final titration step must be at least 2. Using an injection volume of 8 µl and 30 injections, the neutralization point is reached after half of all injections (figure 2). This allows to collect enough data points at saturation and therefore more reliable fitting of the data. If the binding constant is already known, the concentration of the titrand (usually macromolecule in sample cell) can be adjusted to set the *c* value of the current titration inside the recommended experimental window of 5–500. If the binding constant is completely unknown and the stoichiometry *n* equals one, use a molar ratio of 1:10 (titrand to titrant). If the *c* value falls below 5, a ratio of 1:20 is recommended and a sufficient saturation of at least 80 % should be reached at the end of the titration.

**Figure 2:**
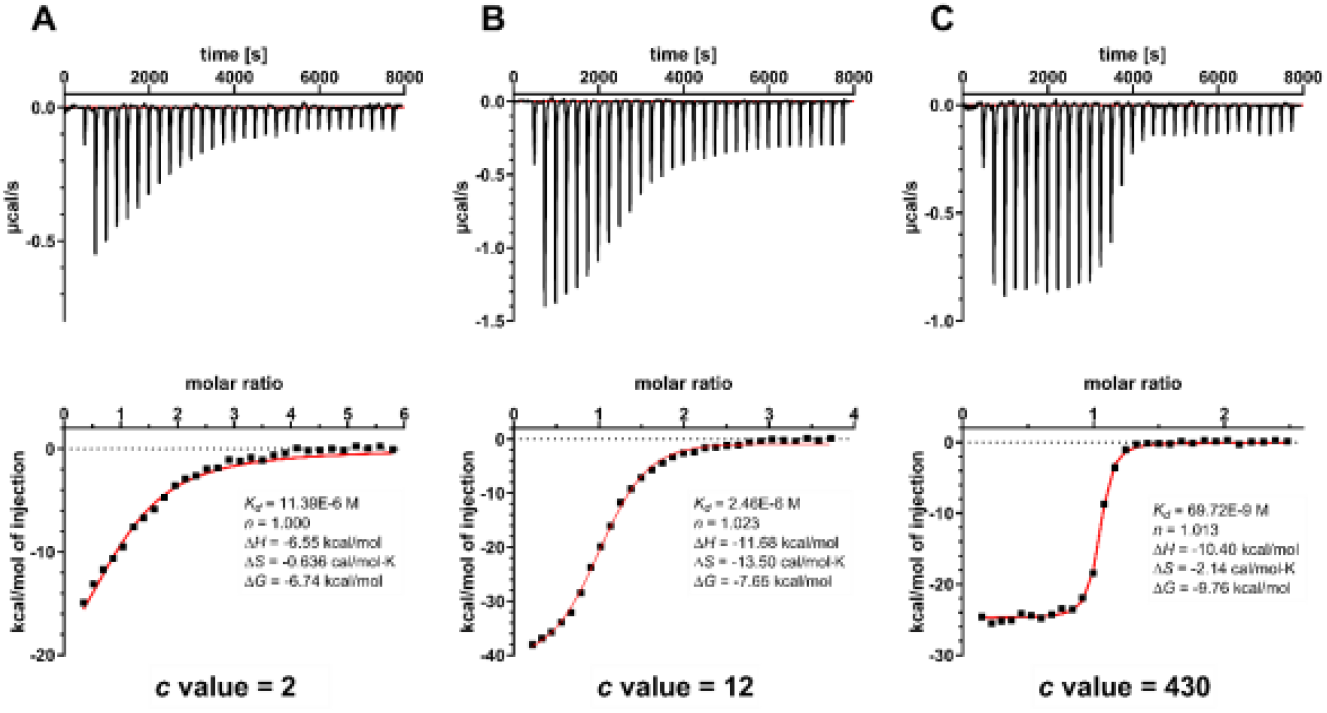
ITC experiments corresponding to the titration of p62-LIR (**a**), LIR28 (**b**) and ANK3 (**c**) against GABARAPL1. The upper panel depicts the thermograms of each binding event. Integrated heat data fitted with a one-site independent binding model along with the calculated thermodynamic parameters is shown below.

The apparent heat recorded over time *Q*_app_ is not only dependent on the heat of ligand binding to the macromolecule, but also on the heats of dilution of the ligand *Q*_dil,L_ and the macromolecule *Q*_dil,M_. Additional nonspecific heat effects *Q*_ns_ could contribute as well. Therefore, the corrected heat is given by:

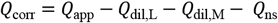

To account for these additive effects, blank titrations should be employed. *Q*_dil,L_ can be assessed by titrating the ligand into buffer without the macromolecule, which usually also incorporates *Q*_ns_. The same applies to *Q*_dil,M_, which values are generally too small to be considered. Due to this, at least one blank titration for the heat of dilution of the titrant is recommended [22]. In order to reduce heat of dilution caused by mixing the buffers of the macromolecule and the ligand, both should be identical in composition. This may either be achieved by dialysis of both components in the same buffer (the dialysate may also be used to dissolve/dilute the ligand) or by multiple runs of buffer exchange with spin filters or buffer exchange columns.

#### 3.2.1 Running the Experiment

The titrations of three peptides, here p62-LIR, LIR28 and the LIR motif of ankyrin 3 (ANK3) will be discussed to provide an overview of how to conduct an incremental titration ITC experiment. LIR28 is an artificial GABARAP selective peptide discovered in a phage display against human Atg8 isoforms [23]. All peptides were titrated into buffer containing isolated GABARAPL1. For further details on the experimental parameters, please refer to Table 1. The herein presented method also applies to measurements of the other 5 homologs and their respective ligands.

**Table 1:** Summary of the experimental parameters employed for the three measurements.

|  | **p62-LIR** | **LIR28** | **ANK3** |
| --- | --- | --- | --- |
| **conc. Peptide [μM]** | 500 | 450 | 240 |
| **conc. Protein [μM]** | 25 | 30 | 30 |
| **Ratio** | 1:20 | 1:15 | 1:8 |
| **Stir Rate** | 350 RPM |  |  |
| **Temperature** | 25 °C |  |  |
| **# Injections, Spacing** | 30, 250 s |  |  |
| **Injection Volume** | 4 μl first, 8 μl others | | |

For sample preparation, the buffer used for all experiments consisted of 50 mM Na_2_HPO_4_ and 100 mM NaCl adjusted to pH 7.0. For proteins containing cysteine residues, such as LC3A or GABARAPL2, 0.5 mM of TCEP should be added. Buffer exchange was conducted via spin columns, with 5 consecutive runs of 10 min at 4 °C and 4000 *g*. For ligands dissolved in DMSO, (here, p62-LIR and LIR28), the same relative amount of DMSO has to be added to the protein sample. Even small deviations in DMSO content between titrant and titrand cause large heats of dilution. Based on previous experiments, the human Atg8 homologs were shown to be stable at DMSO concentrations up to 5 %.

1. To obtain high quality data, samples should be degassed to avoid the formation of bubbles within the cell during the experiment, leading to unpredictable thermal signals.
2. Clean the cell with a suitable cleaning solution based on previous experiments. If a well-behaved system was studied (no apparent precipitation) rinsing with deionized water is sufficient. After aggregation or the use of immiscible organic solvents, it is necessary to employ detergents. Refer to the manufacturer’s instructions for the device. The degassed water within the reference cell also has to be exchanged regularly.
3. Load the sample cell with the appropriate amount of overfill and load the syringe with the titrant as well. To remove any remaining titrant from the syringe, rinse the needle with sample buffer and gently dry the tip using a lint-free tissue. Place the syringe in the sample cell and set the stirring speed to the desired level to allow the system to equilibrate. This is done by monitoring the differential power signal, while waiting for the curve to reach a steady state (some devices support auto-equilibration).
4. Begin the experiment by collecting a baseline of at least 60 seconds. Observe the initial two injections to ensure sufficient spacing, and modify the scheduled injections if necessary.
5. After completion of the titration, remove the syringe and clean the sample cell as mentioned above.

If the initial titration is successful, the blank titration of the ligand into the buffer must be performed identically.

#### 3.2.3 Data Analysis

Determination of *K*_*a*_, Δ*H* and *n* is achieved by nonlinear regression analysis of the experimentally determined data points. The algorithms in use and their implementation might differ, but in general for a one-site binding model, the basis is to find the global minimum of an error surface of *K*_*a*_ and Δ*H*. To yield the apparent heat change for the *i*-th injection Δ*Q*_*i,app*_, the differential power signal for each peak is integrated over time. When the affinity of a ligand to the macromolecule is high, defined by a large *K*_*a*_ or low *K*_*d*_, nearly all the ligand molecules added in the beginning of the titration experiment bind to the macromolecule. This only results in minor changes of Δ*Q*_*i,app*_ for the first injections recorded. As the titration approaches the neutralization point, the absolute heat change decreases until it reaches saturation. At this point, the heat change per injection remains relatively constant. The one-site binding model assumes *n* identical independent binding sites and the *K*_*a*_ can be expressed in terms of Θ, the fraction of sites occupied, with [L] being the free concentration of the ligand:

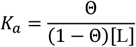

The total ligand concentration [L]_*t*_ can be expressed in terms of the known total concentration of the macromolecule [M]_*t*_:

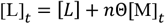

Combining the two prior equations leads to a quadratic equation shown below that can be solved for Θ, which is not directly accessible given that experimental setup:

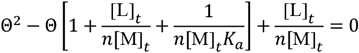

The total heat produced during the titration is given by:

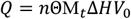

*V*_0_ is the active volume of the sample cell which produces the heat signal. Solving the quadratic equation for Θ and substituting it into the previous equation leads to

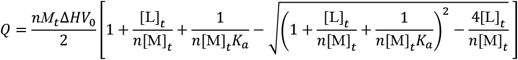

Expressing *Q* in terms of a change in heat per injection Δ*Q*_*i*_ is necessary to then apply the fitting procedure to the model.

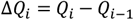

In order to account for the volume being displaced from the sample cell for each injection, corrections are employed for the total concentration of macromolecule and ligand. The model is then employed to fit the data points by initializing the parameters *K*_*a*_, Δ*H* and *n*, calculating Δ*Q*_*i*_ for each injection and then refining them iteratively according to an error function, which describes the quality of the fit given the experimental data [22, 24, 25].

To conclude, after obtaining the experimental data the analysis workflow consists of:

1. Adjusting the baseline to the peaks and defining appropriate windows for the numerical integration of the peaks.
2. The experimental parameters such as employed concentrations, injection volumes and the temperature are either fetched from the files containing the raw data or entered manually.
3. Blank titrations are subtracted to correct the measured *Q*_app_.
4. The correction for the dilution of the samples is usually done automatically within the available analysis software.

Select the appropriate model and fit the data in the case of LC3/GABARAP proteins, usually the one-site binding model. For further information on multiple binding sites, sequential binding or displacement titrations, please refer to the literature [18, 26, 27].

## 4. Notes

- In case of encountering protein aggregation during the experiment, this can be addressed by either altering the buffer system in use or reducing the stirring velocity, while guaranteeing the appropriate mixing of the sample. Crosslinking between proteins, due to disulfide bond formation, may also be a contributing factor which can be resolved through the addition of TCEP or 2-Mercaptoethanol. It is recommended to avoid the use of (2S,3S)-1,4-Bis(sulfanyl)butane-2,3-diol/DTT, because its ring-closing reaction generates significant heats. Another possible solution would be to conduct a reverse titration by adding the protein to the syringe and the low-solubility ligand to the sample cell. Generally speaking, the component of lower solubility should be placed into the sample cell.
- If no well-defined slope is apparent, even though the generated heat is sufficient, this might be due to buffer mismatch between sample cell and syringe material. Ensure that the sample has been prepared properly and every constituent apart from the analytes is equally abundant in both buffers. To further reduce non-specific heats due to friction, keep additives with high viscosity such as glycerol at low concentrations. If the binding event is accompanied by an exchange of protons, the heat of protonation is recorded as well. The number of protons released per binding event is experimentally accessible [28]. 2-Amino-2-(hydroxymethyl)propane-1,3-diol /TRIS for example has a high enthalpy of protonation, by replacing it with another buffer system, heat effects that are due to protonation differences will be reduced.
- In case of a missing saturation at the end of the titration, increase the ratio of the concentrations of ligand to macromolecule. If the rate of Δ*Q*_*i,app*_ decreases too rapidly, it may become necessary to reduce the molar ratio.
- Unexpected discrepancies in stoichiometry may result from imprecise concentration measurements provided as data inputs in the analysis software, solubility limits or significant impurities in protein or ligand samples, partial unfolding of the macromolecule, or selection of an inappropriate binding model.
- The calorimeters and syringes used need to be cleaned thoroughly to reduce artifacts and noisy baselines (especially in experiments in which one of the binding partners precipitated). Refer to the manufacturer’s manual for the compatibility of cleaning agents with the sample cells and the possible cleaning procedures.
- Ensure that the cell volume, injection volume and heat signal are properly calibrated, otherwise, the experimental data may be unreliable [29].

## Acknowledgement

MPS, JD, SK and VVR are grateful for support by the Structural Genomics Consortium (SGC), a registered charity (no: 1097737) that receives funds from Bayer AG, Boehringer Ingelheim, Bristol Myers Squibb, Genentech, Genome Canada through Ontario Genomics Institute, EU/EFPIA/OICR/McGill/KTH/Diamond Innovative Medicines Initiative 2 Joint Undertaking [EUbOPEN grant 875510], Janssen, Merck KGaA, Pfizer and Takeda and by the German Cancer Research Center DKTK and the Frankfurt Cancer Institute (FCI). MPS is funded by the Deutsche Forschungsgemeinschaft (DFG, German Research Foundation), CRC1430 (Project-ID 424228829).

